# *Brucella* effector hijacks endoplasmic reticulum quality control machinery to prevent premature egress

**DOI:** 10.1101/699330

**Authors:** Jean-Baptiste Luizet, Julie Raymond, Thais Lourdes Santos Lacerda, Magali Bonici, Frédérique Lembo, Kévin Willemart, Jean-Paul Borg, Jean-Pierre Gorvel, Suzana P. Salcedo

## Abstract

Perturbation of endoplasmic reticulum (ER) functions can have critical consequences for cellular homeostasis. An elaborate surveillance system known as ER quality control (ERQC) ensures that only correctly assembled proteins reach their destination. Persistence of misfolded or improperly matured proteins upregulates the unfolded protein response (UPR) to cope with stress, activates ER associated degradation (ERAD) for delivery to proteasomes for degradation. We have identified a *Brucella abortus* type IV secretion system effector called BspL that targets Herp, a key component of ERQC and is able to augment ERAD. Modulation of ERQC by BspL results in tight control of the kinetics of autophagic *Brucella*-containing vacuole formation, preventing premature bacterial egress from infected cells. This study highlights how bacterial pathogens may hijack ERAD components for fine regulation of their intracellular trafficking.

## Introduction

The endoplasmic reticulum (ER) is the largest organelle in the cell and plays numerous functions vital for maintaining cellular homeostasis. It is the major site for protein synthesis of both secreted and integral membrane proteins as well as exporting of newly synthesised proteins to other cellular organelles. Disturbance or saturation of the folding-capacity of the ER leads to a complex stress response that has evolved to help cells recover homeostasis or, if necessary, commit them to death. The ER relies on a complex surveillance system known as ER quality control (ERQC) that ensures handling of misfolded, misassembled or metabolically regulated proteins (Braakman and Bulleid, 2011). Once retained in the ER, these proteins are retrotranslocated back into the cytosol to be ubiquitinated and degraded by the proteasome, a process known as ER-associated degradation (ERAD) (Wu and Rapoport, 2018). Alternatively, ERAD-resistant proteins can be degraded *via* ERQC-autophagy (Houck et al., 2014).

In response to ER perturbations, particularly following the accumulation of toxic amounts of misfolded proteins, ER stress ensues and cells activate a set of inter-connected pathways that are collectively referred to as the unfolded protein response (UPR) that have a critical role in restoring homeostasis (Walter and Ron, 2011). The UPR is regulated by three ER membrane sensors, the inositol-requiring enzyme I (IRE1), double-stranded RNA-activated protein kinase-like ER kinase (PERK) and activating transcription factor 6 (ATF6). In non-stress conditions these are kept inactive thanks to their association with the ER chaperone BiP. Upon stress, BiP is dislodged from the luminal domains of the three sensors which leads to their activation and induction of specialized transcriptional programs. The IRE1 and ATF6 pathways are involved in induction of the transcription of genes encoding for protein-folding chaperones and ERAD-associated proteins (Hetz and Papa, 2018). Whereas PERK sensing is particularly important in control of autophagy, protein secretion and apoptosis (Hetz and Papa, 2018).

The homocysteine-inducible ER stress protein (Herp) is an ER membrane protein that is highly upregulated during ER stress by all UPR branches (Kokame et al., 2000; Ma and Hendershot, 2004). Herp is a key component of ERQC that plays a protective role in ER stress conditions (Chan et al., 2004; Tuvia et al., 2007). It is an integral part of the ERAD pathway, enhancing the protein loading and folding capacities of the ER. In addition, it acts as a hub for membrane association of ERAD machinery components, stabilizing their interactions with substrates at ERQC sites (Leitman et al., 2014) and facilitating their retrotranslocation (Huang et al., 2014). Furthermore, as Herp is also in a complex with the proteasome it may aid delivery of specific retrotranslocated substrates to the proteasome for degradation (Kny et al., 2011; Okuda-Shimizu and Hendershot, 2007).

Given its importance for cellular homeostasis, the ERQC represents a prime target for microbial pathogens. Indeed, a growing number of bacterial pathogens have been shown to hijack ERQC pathways, especially by modulating UPR (Celli and Tsolis, 2014). For example, *Legionella pneumophila* secretes several effector proteins that repress CHOP, BiP and XBP1s at the translational level, resulting in UPR inhibition and decrease in inflammation (Hempstead and Isberg, 2015). Another pathogen for which modulation of UPR plays a critical role during infection is *Brucella* spp., a facultative intracellular pathogen that causes brucellosis, a zoonosis still prevalent worldwide. *Brucella abortus* has been shown to induce UPR (de Jong et al., 2012; Smith et al., 2013), and more specifically the IRE1 pathway, contributing to enhanced inflammation, a process particularly relevant in the context of colonization of the placenta and abortion (Keestra-Gounder et al., 2016). However, activation of IRE1 is also important for *Brucella* trafficking and subsequent *Brucella* multiplication (Qin et al., 2008; Smith et al., 2013). After cellular uptake, *Brucella* is found in a membrane bound compartment designated endosomal *Brucella-*containing vacuole (eBCV) which transiently interacts with early and late endosomes, undergoing limited fusion with lysosomes (Starr et al., 2008). Bacterial are then able to sustain interactions with ER exit sites (ERES) a process that requires the activity of the small GTPases Sar1 (Celli et al., 2005) and Rab2 (Fugier et al., 2009) and results in the establishment of an ER-derived compartment suited for multiplication (replicative or rBCV). UPR induction by *Brucella* is necessary for this trafficking step, as the formation of rBVCs is dependent on IRE1 activation by the ERES-localized protein Yip1A, which mediates IRE1 phosphorylation and dimerization (Taguchi et al., 2015). Once rBCVs are established, *Brucella* is capable of extensive intracellular replication, without induction of cell death. Instead, at late stages of the intracellular cycle, rBCVs reorganize and fuse to form large autophagic vacuoles (aBCVs) that will mediate bacterial exit from infected cells (Starr et al., 2011). The bacterial factors behind the switch between rBCVs and aBCVs remain uncharacterized.

*Brucella* relies on a type 4 secretion system (T4SS), encoded by the *virB* operon and induced during eBCV trafficking to translocate bacterial effectors into host cells and directly modulate cellular functions. However, only a few effectors have been characterized and for which we have a full grasp of how they contribute towards pathogenesis. This system has been implicated in the induction of UPR during infection and a subset of these effectors has been shown to modulate ER-associated functions. VceC interacts with the ER chaperone BiP to activate the IRE1 pathway, which results in NOD1/NOD2 activation and up-regulation of inflammatory responses (de Jong et al., 2012; Keestra-Gounder et al., 2016). BspA, BspB and BspF have all been implicated in blocking of ER secretion (Myeni et al., 2013). In particular, BspB was shown to interact with the conserved oligomeric Golgi (COG) complex to redirect vesicular trafficking towards the rBCVs (Miller et al., 2017). Several other effectors that localize in the ER when ectopically expressed have been shown to induce UPR or control ER secretion, but the mechanisms involved remain uncharacterized.

In this study, we identify a new T4SS effector of *Brucella abortus*, that we designate as *Brucella-*secreted protein L (BspL) that targets a component of the ERAD machinery, Herp. BspL enhances ERAD and delays the formation of aBCVs, preventing early bacterial release from infected cells which helps maintain cell to cell spread efficiency.

## Results

### BspL is a *Brucella* T4SS effector protein

Bacterial effectors are often similar to eukaryotic proteins or contain domains and motifs that are characteristic of eukaryotic proteins. Multiple bacterial effectors benefit from the host lipidation machinery for targeting eukaryotic membranes. Some of these contain a carboxyl-terminal CAAX tetrapeptide motif (C corresponds to cysteine, A to aliphatic amino acids and X to any amino acid) that serves as a site for multiple post-translation modifications and addition of a lipid group which facilitates membrane attachment, such as SifA from *Salmonella enterica* (Boucrot et al., 2003; Reinicke, 2005) and AnkB from *Legionella pneumophila* (Price et al., 2010). Previous work highlighted several *Brucella* encoded proteins that contain putative CAAX motifs (Price et al., 2010) which could therefore be T4SS effectors. In this study, we focused on one of these proteins encoded by the gene BAB1_1533 (YP_414899.1), that we have designated BspL for *Brucella*-secreted protein L. We first determined if BspL was translocated into host cells during infection. We constructed a strain expressing BspL fused to the C-terminus of the TEM1 ß-lactamase (encoded by *bla*) and infected RAW macrophage-like cells for different time-points. A Flag tag was also included for control of protein expression. The fluorescent substrate CCF2 was added and the presence of fluorescent emission of coumarin, resulting from cleavage by the cytosolic TEM1 lactamase, was detected by confocal microscopy. This assay is widely used in the *Brucella* field and we included the T4SS effector VceC as a positive control (de Jong et al., 2008), which showed the highest level of secretion at 24h post-infection in our experimental conditions (Figure 1A). We found that TEM1-BspL was secreted into host cells as early as 4h post-infection, with a slight peak at 12h post-infection, CCF2 cleavage was still detected at 24h post-infection (Figure 1A). This phenotype was fully dependent on the T4SS as a Δ*virB9* mutant strain did not show any coumarin fluorescence (Figure 1A and B). This was not due to lack of expression of TEM1-BspL as both the wild-type and the Δ*virB9* strains carrying the *bla*::*bspL* plasmid showed equivalent levels of TEM1-BspL expression (Figure 1C). Together, these results show BspL is a T4SS effector.

**Figure 1.**
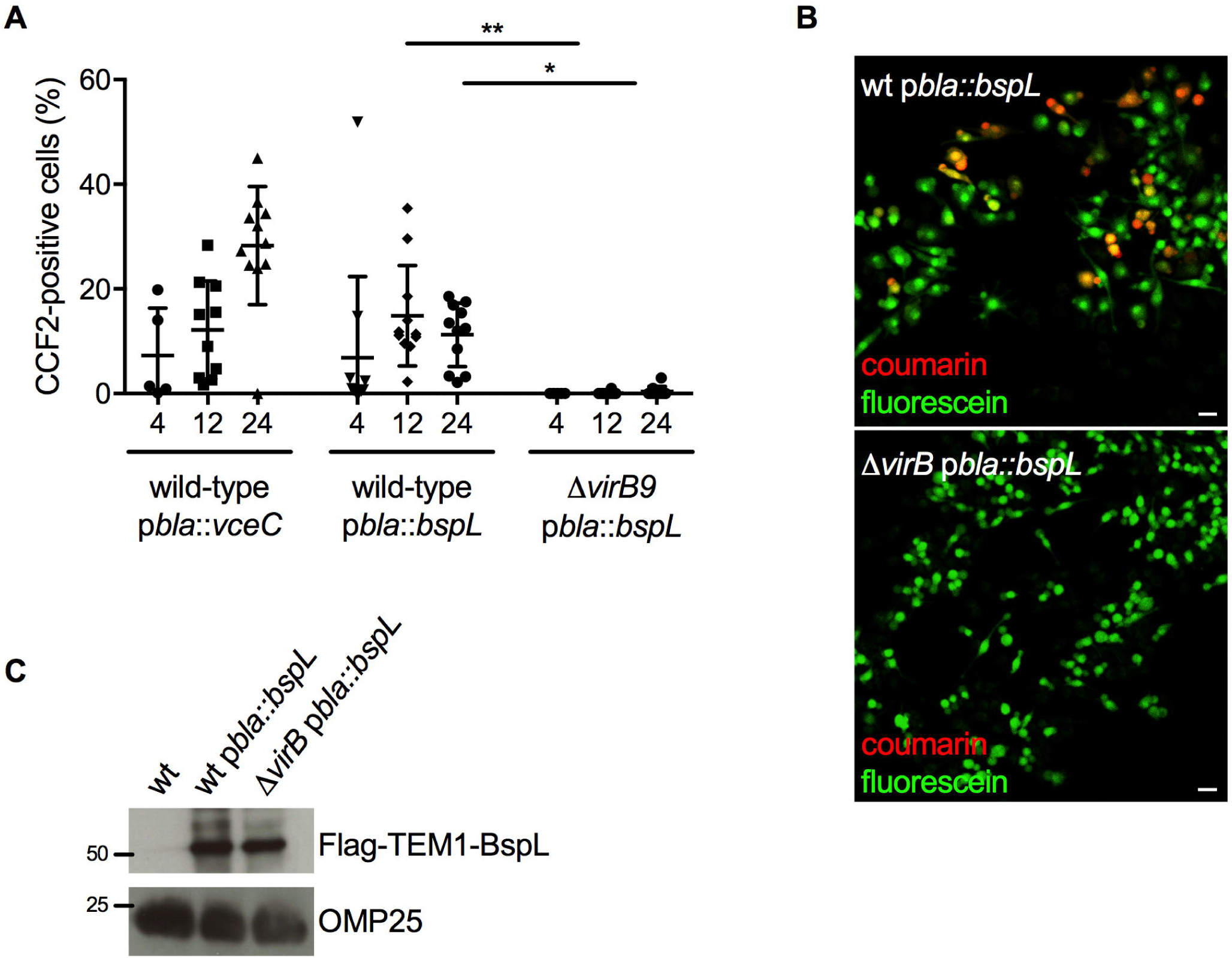
BspL is a T4SS effector translocated into host cells during *B. abortus* infection. (A) Macrophage-like cell line (RAW) was infected with *B. abortus* carrying a plasmid encoding for *bla* fused with BspL (p*bla::bspL*) to enable expression of TEM-BspL. Cells were infected with either wild-type *B. abortus* or Δ*virB9* carrying this plasmid. A positive control of wild-type expressing *bla::vceC* was included. At 4, 12 or 24h post-infection, cells were incubated with fluorescent substrate CCF2-AM, fixed and the percentage of cells with coumarin emission quantified using an automated plugin. More than a 1000 cells were quantified for each condition from 3 independent experiments and data represent means ± standard deviations. Kruskal-Wallis with Dunn’s multiple comparisons test was used and P = 0.0019 between wild-type p*bla::bspL* and Δ*virB9* p*bla::bspL* at 12h (**) and 0.171 at 24h (*). Not all statistical comparisons are shown. (B) Representative images of cells infected for 24h with *B. abortus* wild-type or Δ*virB9* carrying p*bla::bspL*. Cells were incubated with CCF2 and the presence of translocated TEM1-BspL detected by fluorescence emission of coumarin (red). Scale bars correspond to 5 µm. (C) The expression of TEM1-BspL in the inocula of wild-type and Δ*virB9* strains was controlled by western blotting thanks to the presence of a FLAG tag in the construct. The membrane was probed with an anti-Flag antibody (top) or anti-Omp25 (bottom) as a loading control. A sample from wild-type without the plasmid was included as a negative control. Molecular weights are indicated (KDa).

### Ectopically expressed BspL accumulates in the ER, does not interfere with host protein secretion but induces the UPR

BspL is very well conserved in the *Brucella* genus, it is 170 amino acids long (Figure S1A) and is approximately 19 kDa. BspL does not share any homology to eukaryotic proteins nor to other bacterial effectors. Its nucleotide sequence encodes for a sec secretion signal, a feature commonly found in other *Brucella* effectors (Marchesini et al., 2011). In addition, it contains a hydrophobic region that may constitute a transmembrane domain as well as a proline rich region, with seven consecutive prolines that may be relevant in interactions with eukaryotic proteins. To gain insight into the function of BspL we ectopically expressed HA, myc or GFP-tagged BspL in HeLa cells. We found BspL accumulated in the ER, as can be seen by the co-localization with calnexin (Figure 2A and S1B, S1C), an ER membrane protein and chaperone. Unlike what has been reported for VceC (de Jong et al., 2012), the structure of the ER remained relatively intact upon BspL expression. Deletion of the C-terminal tetrapeptide sequence, which could correspond to a potential lipidation motif had no effect on the ER localization of BspL in transfection (Figure S1B, bottom panel), as it significantly overlapped with the full-length protein when co-expressed in the same cell (Figure S1C).

**Figure 2.**
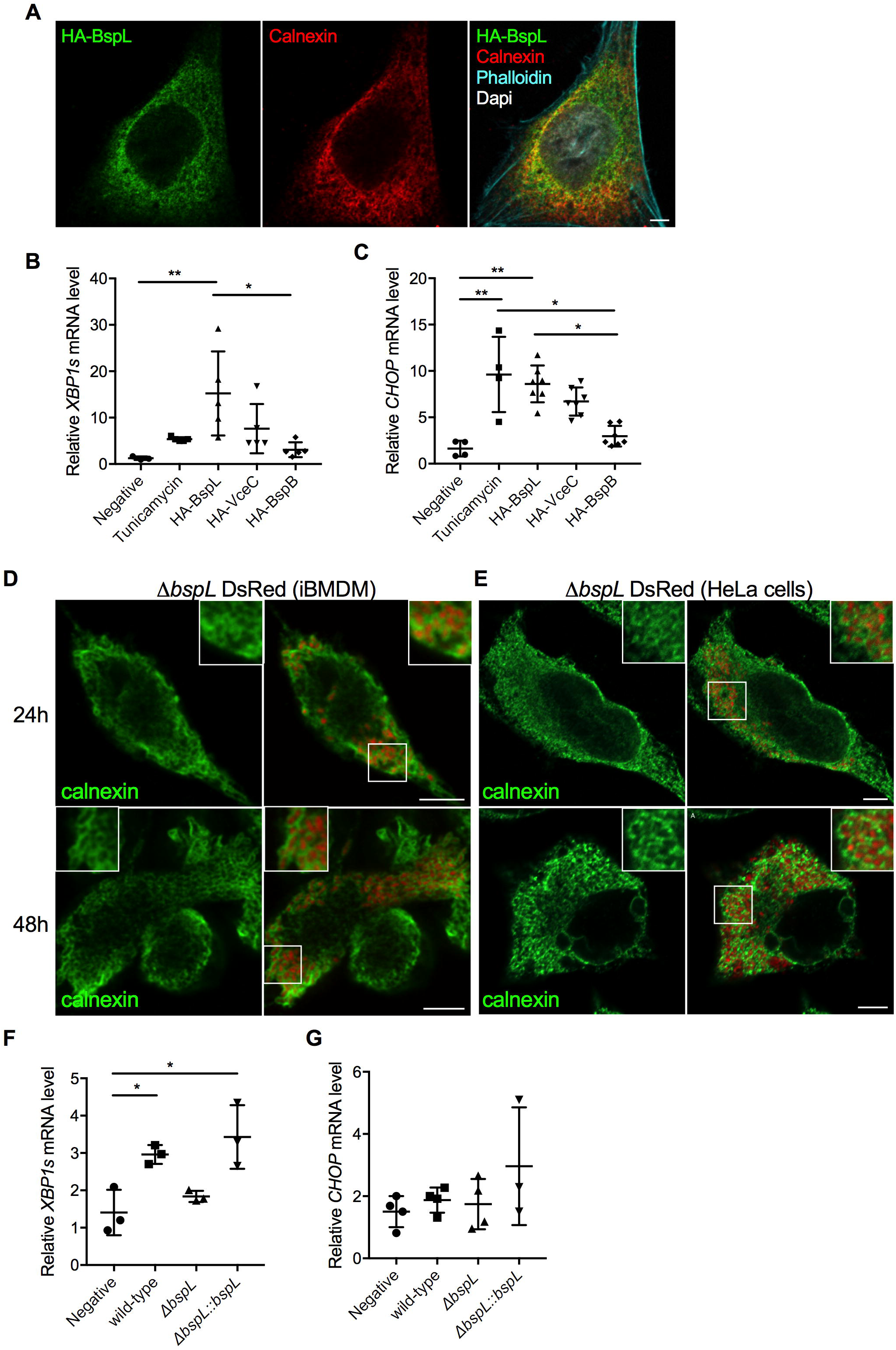
BspL does not impact early BCV trafficking but contributes to UPR induction at late stages of the infection. (A) Confocal microscopy image showing the intracellular localization of HA-BspL expressed in HeLa cells labelled with an anti-HA antibody (green) and ER marker calnexin (red). Phalloidin (cyan) was used to label the actin cytoskeleton and Dapi (white) for the nucleus. (B) Quantification of mRNA levels of *XBP1s* and (C) *CHOP* by quantitative RT-PCR obtained from HeLa cells expressing HA-BspL, HA-VceC or HA-BspB for 24h. Cells transfected with empty vector pcDNA3.1 were included as a negative control and cells treated tunicamycin at 1µg/µl for 6h as a positive control. Data correspond to the fold increase in relation to an internal control with non-transfected cells. Data are presented as means ± standard deviations from at least 4 independent experiments. Kruskal-Wallis with Dunn’s multiple comparisons test was used and P = 0.042 between negative and HA-BspL (**) and 0.0383 between HA-BspL and HA-BspB (*) for *XBP1s*. For *CHOP*, P = 0.0184 between negative and tunicamycin (*); 0.0088 between negative and HA-BspL (**); 0.0297 bteween tunicamycin and HA-BspB (*) and 0.011 between HA-BspL and HA-BspB (*). All other comparisons ranked non-significant. (D) Representative images of rBCVs from Δ*bspL-*expressing DSred infected iBMDM or (E) HeLa cells at 24 and 48h post-infection, labelled for calnexin (green). (F) Quantification of mRNA levels of *XBP1s* and (G) *CHOP* by quantitative RT-PCR obtained from iBMDMs infected with wild-type, Δ*bspL* or the complemented Δ*bspL::bspL* strains for 48h. Mock-infected cells were included as a negative control. Data correspond to the fold increase in relation to an internal control with non-infected cells. Data are presented as means ± standard deviations from at least 3 independent experiments. Kruskal-Wallis with Dunn’s multiple comparisons test was used and, for *XBP1s*, P = 0.042 between negative and HA-BspL (**) and 0.0352 between the negative control and wild-type infected cells (*) and 0.0111 between negative and the complemented Δ*bspL::bspL* infected cells (*). All other comparisons ranked non-significant with this test.

Our observations suggest BspL is part of a growing number of *Brucella* effectors that accumulate in the ER when ectopically expressed, including VceC, BspB and BspD (de Jong et al., 2012; Myeni et al., 2013). We therefore investigated if BspL shared any of the ER modulatory functions described for other effectors, notably interference with ER secretion as BspB (Miller et al., 2017; Myeni et al., 2013) or induction of ER stress as VceC (de Jong et al., 2012; Keestra-Gounder et al., 2016).

To determine the impact of BspL on host protein secretion we used the secreted embryonic alkaline phosphatase (SEAP) as a reporter system. HEK cells were co-transfected with the vector encoding SEAP and vectors encoding different *Brucella* effectors. We chose to work with HA-BspL, to allow direct comparison with previously published HA-BspB that blocks ER secretion and HA-BspD as a negative control (Myeni et al., 2013). Expression of the GDP-locked allele of the small GTPase Arf1[T31N], known to block the early secretory pathway, was used as a control for efficient inhibition of secretion (Figure S1D). As previously reported, we found that expression of HA-BspB drastically reduced SEAP secretion (Figure S1D). In contrast, HA-BspL did not impact SEAP secretion to the same extent as BspB, having an effect equivalent to HA-BspD previously reported not to affect host protein secretion (Myeni et al., 2013).

We next investigated whether ER targeting of BspL was accompanied with activation of the UPR, an important feature of *Brucella* pathogenesis. In the case of *B. abortus*, IRE1 is the main pathway activated (de Jong et al., 2012) which leads to splicing of the mRNA encoding the transcription factor X-box-binding protein 1 (XBP1) which in turn induces the expression of many ER chaperones and protein-folding enzymes. The second branch of the UPR dependent on PERK may also be of relevance in *Brucella* infection (Smith et al., 2013). Under prolonged stress conditions, this UPR branch leads to the up-regulation of the transcription factor C/EBP-homologous protein (CHOP) which induces expression of genes involved apoptosis. We therefore monitored *XBP1s* and *CHOP* transcript levels following ectopic expression of HA-BspL, in comparison to HA-VceC, established as an ER stress inducer and HA-BspB, known not to induce ER stress. Treatment with tunicamycin, a chemical ER stress inducer was also included. We found that over-expression of HA-BspL induced an increase of both *XBP1s* and *CHOP* transcription, to levels even higher than HA-VceC (Figure 2B and C). These results suggest BspL may induce ER stress.

### BspL is not involved in establishment of an ER-derived replication niche but is implicated in induction of ER stress during infection

As UPR has been implicated in the establishment of rBCVs (Taguchi et al., 2015) and intracellular replication (Qin et al., 2008; Smith et al., 2013; Taguchi et al., 2015) of *Brucella* we next investigated the intracellular fate of a *B. abortus* 2308 strain deleted for *bspL* in comparison with the wild-type. Two cellular models were used, HeLa cells and an immortalized cell line of bone marrow-derived macrophages (iBMDM). We found that the Δ*bspL* strain replicated as efficiently as the wild-type in both iBMDM (Figure S2A) and HeLa cells (Figure S2B). In terms of intracellular trafficking no obvious differences were observed in the establishment of rBCVs at 24 and 48h post-infection, as Δ*bspL* BCVs were nicely decorated with the ER marker calnexin in both cell types (Figure 2D and E) as observed for the wild-type strain (Figure S2C and D). As this is the first report to our knowledge to use iBMDM in *Brucella* infections, we confirmed this observation by quantifying the percentage of BCVs positive for calnexin and the lysosomal associated membrane protein 1 (LAMP1) in comparison with the wild-type at 24 and 48 post-infection (Figure S2E and F, respectively). The wild-type strain in this cellular model behaved as expected forming the typical rBCVs.

As in transfected cells we found that BspL induced UPR, we next monitored the levels of *XBP1s* and *CHOP* transcripts during infection. Since the rate of infected cells is too low to detect ER stress in HeLa cells, these experiments were only performed in iBMDMs. As expected, the wild-type *B. abortus* strain induced an increase in the levels of transcription of *XBP1s* in relation to the mock-infected control iBMDM at 48h post-infection (Figure 2F). In contrast, Δ*bspL* infected macrophages showed decreased *XBP1s* transcript levels compared to the wild-type (Figure 2G). Furthermore, the wild-type phenotype could be fully restored by expressing a chromosomal copy of *bspL* in the Δ*bspL* strain, confirming that BspL specifically contributes towards induction of the IRE1 branch of the UPR during infection (Figure 2F). We did not observe an increase in *CHOP* transcript levels in iBMDM infected with the wild-type nor Δ*bspL* strains in comparison to the mock-infected cells (Figure 2G), suggesting that *B. abortus* does not significantly induce the PERK-dependent branch of the UPR at this stage of the infection.

### BspL interacts with Herp, a key component of ERQC

To gain insight into the function of BspL we set out to identify its interacting partners. A yeast two-hybrid screen identified 7 candidates: eukaryotic translation initiation factor 4A2 (EIF4A2), pyruvate dehydrogenase beta (PDHB), MTR 5-methyltetrahydrofolate-homocysteine methyltransferase, Bcl2-associated athanogene 6 (BAG6), ARMCX3 armadillo repeat containing protein (Alex3), homocysteine-inducible ER protein with ubiquitin like domain (Herpud or Herp) and Ubiquilin2 (Ubqln2).

In view of our previous results for BspL showing ER localization and induction of UPR we decided to focus on Alex3, Herp and Ubiquilin2 which are rarely present or even absent in the database of false positives for this type of screen (http://crapome.org/). Alex3 is a mitochondrial outer membrane protein that has been implicated in regulation of mitochondrial trafficking (Serrat et al., 2013). As ER and mitochondria extensively interact, Alex3 could constitute an interesting target. Herp is an ER membrane protein playing a role in both the UPR and the ERAD system whereas Ubiquilin2 is implicated in both the proteasome and ERAD and, interestingly, shown to interact with Herp (Kim et al., 2008). In view of these different targets we decided to carry out an endogenous co-immunoprecipitation in cells expressing HA-BspL. As controls for detecting non-specific binding, we also performed co-immunoprecipitations from cells expressing two other ER-targeting effectors, HA-BspB and HA-VceC. We then probed the eluted samples with antibodies against Alex3, Ubiquilin2 or Herp to detect if any interactions could be observed. We found that Alex3 was co-immunoprecipitated with all 3 effectors suggesting a potentially non-specific interaction with the effectors or the resin itself (Figure 3A). In contrast, no interactions were observed with Ubiquilin2, which was detected only in the flow through fractions. However, we found that endogenous Herp specifically co-immunoprecipated with HA-BspL and not the other effectors (Figure 3A), suggesting Herp and BspL form a complex within host cells. Taken together with the yeast two-hybrid data, we can conclude that BspL directly interacts with Herp. Consistently, over-expressed BspL co-localized with Herp by microscopy (Figure 3B).

**Figure 3.**
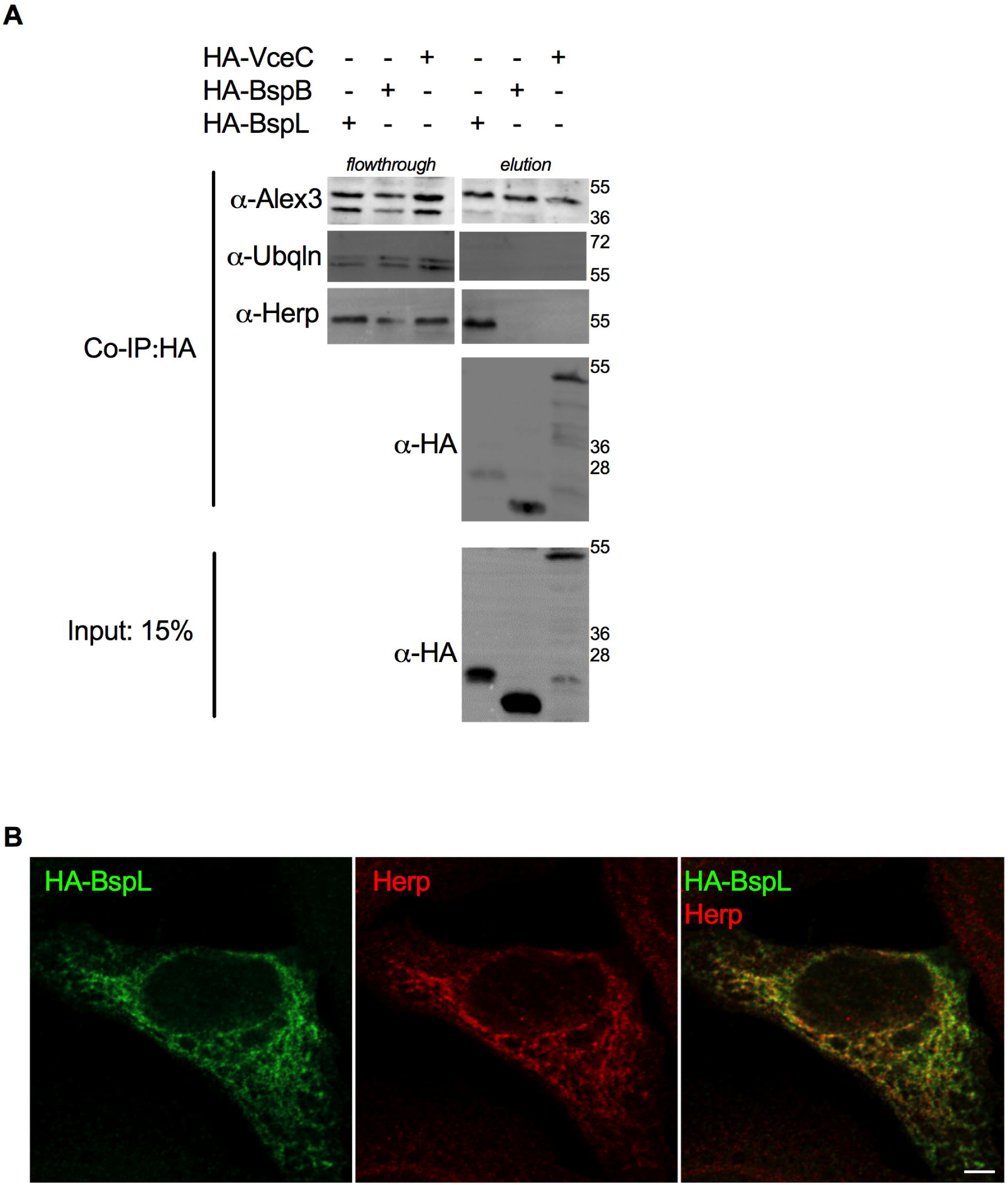
BspL specifically interacts with the ERAD component Herp. (A) Co-immunoprecipitation (co-IP) from cell extracts expressing either HA-BspL, HA-BspB and HA-VceC using HA-trapping beads. Flow through and elutions were probed with antibodies against Alex3, Ubiquilin (Ubqln) and Herp in succession. The level of each effector bound to the beads was revealed with an anti-HA antibody and 15% of the input used for the co-IP shown (at the bottom). Molecular weights are indicated (KDa). (B) Representative confocal micrograph of HeLa cells expressing HA-BspL (green) and labelled for Herp (red). Scale bar corresponds to 5 µm.

### BspL facilitates degradation of TCR**α** *via* ERAD independently of ER stress

Herp is a key component of ERAD, strongly up-regulated upon ER stress. Indeed, during *B. abortus* infection we observed an up-regulation of *HERP* transcripts (Figure S3A), consistent with *XBP1s* induction, although these differences were not statistically significant with the number of replicates carried out. However, inhibition of Herp using siRNA (Figure S3B) showed that ER stress induced following ectopic expression of BspL was not dependent on Herp (Figure S3C and D), suggesting BspL interaction with Herp is mediating other functions in the cell.

Therefore, we next investigated if BspL could directly impact ERAD. We used expression of T cell receptor alpha (TCRα) as reporter system, as this type I transmembrane glycoprotein has been shown to be a canonical ERAD substrate, quickly degraded (Feige and Hendershot, 2013; Lippincott-Schwartz et al., 1988). TCRα is transferred across the ER membrane, where is becomes glycosylated and fails to assemble. This in turn induces its retrotranslocation back to the cytosol to be degraded by the proteasome. Cycloheximide treatment for 4 h was used to block protein synthesis, preventing replenishment of TCR pools and allowing for visualization of ERAD-mediated degradation of TCRα. When HEK-293T cells, which do not naturally express TCR were transfected with HA-TCRα and treated with cycloheximide, a decrease in HA-TCRα was observed, indicative of degradation (Figure 4A, red arrow). Strikingly, expression of BspL induced very strong degradation of TCRα (Figure 4A). This is accompanied by the appearance of a faster migrating band at around 25 KDa (blue arrow), that nearly disappears upon cycloheximide treatment suggesting this TCRα peptide is efficiently degraded by the proteasome. It is important to note that the 25 KDa band is also present when HA-TCRα is expressed alone (lane 2 of Figure 4A, blue arrow) suggesting it is a natural intermediate of HA-TCRα degradation.

**Figure 4.**
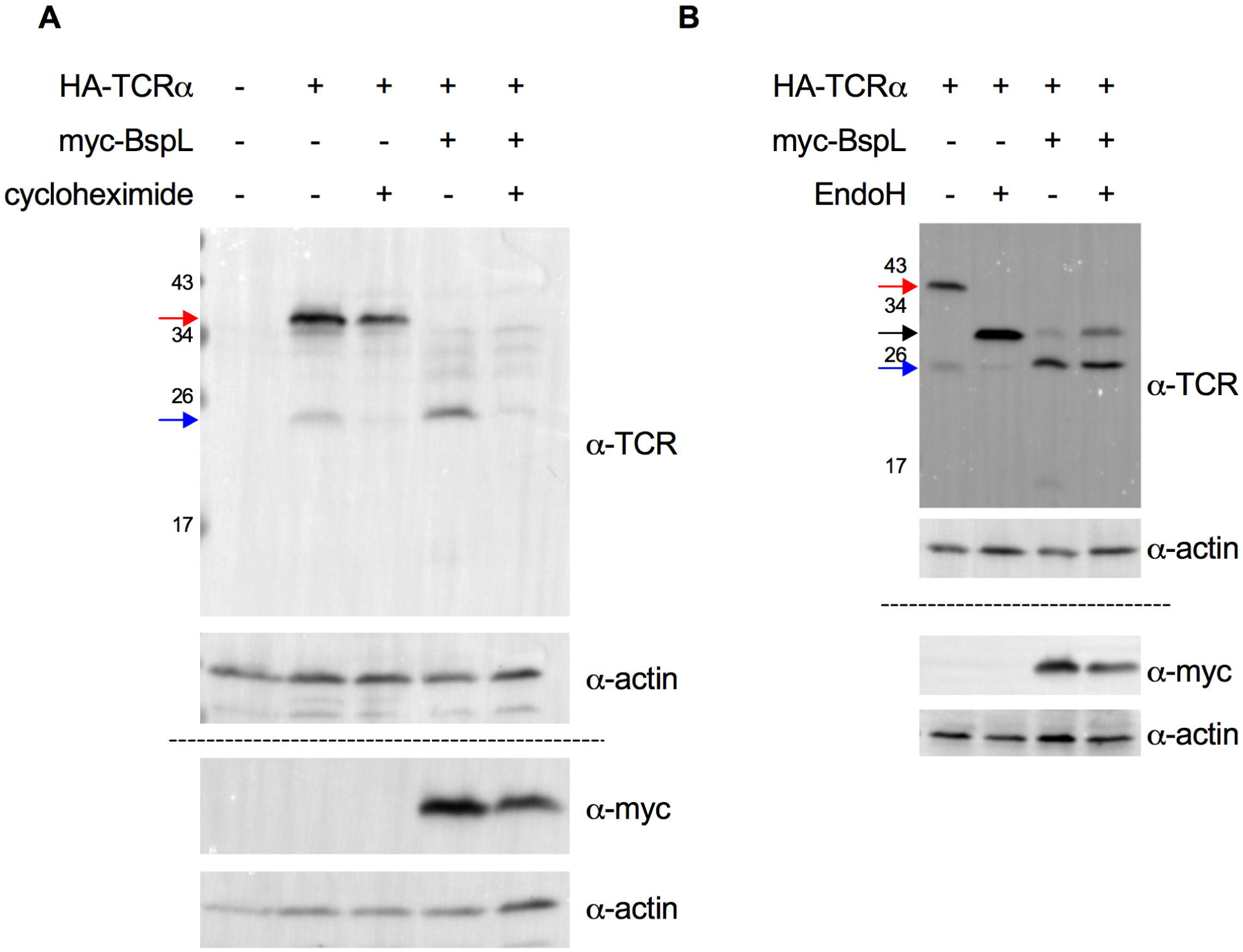
BspL enhances ERAD degradation of TCRα. (A) HEK 293T cells were transfected with HA-TCRα in the absence or presence of myc-BspL for 24h. Where indicated, cells were treated with 50 µg/ml cycloheximide for the last 4h. The blot was probed first with an anti-TCR antibody followed by anti-actin. The same samples were loaded onto a separate gel (separated by dashed line) for probing with an anti-myc and anti-actin to confirm the expression of myc-BspL. Molecular weights are indicated (KDa) and relevant bands described in the text highlighted with different coloured arrows. (B) HEK 293T cells were transfected with HA-TCRα in the absence or presence of myc-BspL for 24h and samples treated with EndoH where indicated. The blot was probed first with an anti-TCR antibody followed by anti-actin. The same samples were loaded onto a separate gel (separated by dashed line) for probing with an anti-myc and anti-actin to confirm the expression of myc-BspL. Molecular weights are indicated (KDa) and relevant bands described in the text highlighted with different coloured arrows.

To determine if the enhanced effect of BspL on TCRα degradation is a side-effect of ER stress, cells were treated with TUDCA which strongly inhibited both *XBP1s* and *CHOP* transcript levels induced by either tunicamycin, BspL or VceC (Figure S3E and F). In the presence of TUDCA, BspL was still found to enhance HA-TCRα degradation showing this is occurring in an ER stress-independent manner (Figure S4).

As the TCRα subunit undergoes N-glycosylation in the ER, we wondered if the faster migrating band of TCRα induced by BspL corresponded to non-glycosylated form of TCRα. We therefore treated samples with EndoH, which deglycosylates peptides. Upon EndoH treatment we observed deglycosylated HA-TCRα (second lane, Figure 4B, black arrow), confirming the reporter system is being processed normally. In the BspL expressing samples (lanes 3 and 4, Figure 4B), a slight band corresponding to the non-glycosylated TCRα could also be detected particularly after EndoH treatment, confirming that BspL does not prevent TCRα from entering the ER and being glycosylated. The dominant TCRα band induced upon BspL expression (around 25 KDa, blue arrow) migrates faster than the non-glycosylated form resulting from EndoH treatment (black arrow) and does not appear to be sensitive to EndoH. This may therefore correspond to a natural truncated non-glycosylated form of HA-TCRα. Consistently, this band is also present in the absence of BspL (lane 1, Figure 4B, blue arrow). Together these data indicate that BspL is a strong inducer of ERAD.

### ERAD is required for different stages of intracellular lifecycle of *Brucella*

The role of ERAD in the *Brucella* intracellular life cycle has not yet been investigated to our knowledge. We therefore decided to block ERAD using eeyarestatin, an established inhibitor of this system. Unfortunately, prolonged treatment at the concentration necessary for full inhibition of ERAD induced detachment of infected iBMDM. Nonetheless, we were able to carry out this experiment in HeLa cells, which showed significant resistance to the eeyarestatin treatment. Total CFU counts after addition of eeyarestatin at 2h post-infection showed a significant decrease in bacterial counts at 48h, suggesting a potential inhibition of replication (Figure 5A). However, microscopy observation of infected cells at this time-point clearly showed extensive replication of bacteria even in the presence of eeyarestatin (Figure 5B), suggesting that the drop of CFU observed was a result of exit of bacteria from infected cells rather than inhibition of intracellular replication. Consistently, we observed significant numbers of extracellular bacteria as well as many cells infected with only a few bacteria potentially resulting from re-infection. These results suggest that blocking of ERAD during early stages of infection would favour intracellular replication. To confirm this possibility, we counted by microscopy the number of bacteria per cell at 24h post-infection and indeed found a higher replication rate upon eeyarestatin treatment (Figure 5C). We therefore hypothesized that *Brucella* might block ERAD during early stages of the infection to favour establishment of an early replication niche, a phenotype clearly not dependent on BspL, as we have shown it is not implicated in the establishment of rBCVs and when ectopically expressed it induces ERAD. We therefore, wondered if BspL could intervene at a later stage of the infection to induce ERAD *via* its interaction with Herp.

**Figure 5.**
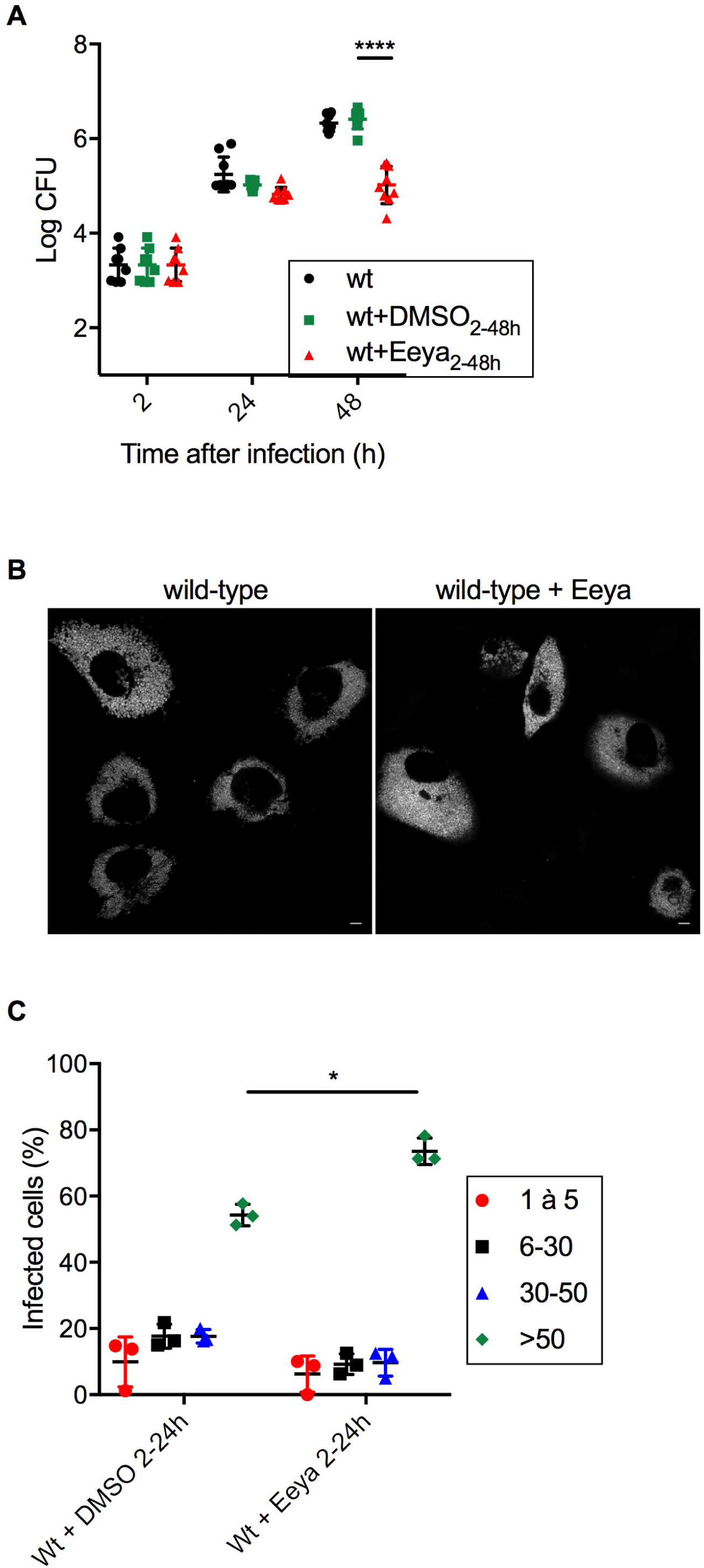
Blocking of ERAD at early stages of the infection enhances intracellular replication and accelerates bacterial release. (A) Bacterial counts (CFU) at 2, 24 and 48h post-infection with either the wild-type without any treatment (wt, black) or in the presence of 8 µM eeyarestatin (wt+Eeya, red) or the equivalent amount of DMSO (wt+DMSO, green). Data correspond to means ± standard deviations from 6 independent experiments. A two-way ANOVA was used yielding a P < 0.0001 (****) between wild-type+DMSO with wild-type+Eeya at 48h. Other comparisons are not indicated. (B) Representative confocal images of HeLa cells infected with the wild-type DSRed or following treatment eeyarestatin at 48h post-infection. (C) Microscopy bacterial counts at 24h post-infection with either the wild-type with DMSO or in the presence of 8 µM eeyarestatin. Data is presented as the percentage of cells containing 1 to 5 bacteria per cell (red), 6 to 30 (black), 30 to 40 (blue) or more than 50 (green). Data correspond to means ± standard deviations from 3 independent experiments. A two-way ANOVA test was used yielding a P= 0.0003 (***) between wild-type+DMSO with wild-type+Eeya at 48h. Other comparisons are not indicated.

### BspL delays premature bacterial egress from infected cells

The late stage of the intracellular cycle of *Brucella* relies on induction of specific autophagy proteins to enable the formation of aBCVs characterized as large vacuoles with multiple bacteria decorated with LAMP1 (Starr et al., 2011). In our experimental conditions aBCVs could be clearly observed in iBMDM infected for 65h with wild-type *B. abortus* (Figure 6A). We therefore investigated if BspL was involved in formation of aBCVs. Strikingly, Δ*bspL* aBCVs could be detected as early as 24h, with nearly 30% of infected cells showing aBCVs at 48h post-infection compared to less than 10% for wild-type infected cells (Figure 6B and C). Importantly, complementation of the Δ*bspL* strain fully restored the wild-type phenotype. These results strongly suggest that BspL is involved in delaying the formation of aBCVs during *B. abortus* macrophage infection. Consistently, imaging of Δ*bspL* infected iBMDM at 48h, revealed the presence of high numbers of extracellular bacteria as well as cells with single bacteria or a single aBCV (Figure 6D), suggestive of re-infection and reminiscent of what was observed following eeyarestatin treatment that blocks the ERAD. In contrast, wild-type infected iBMDM at the same time-point showed none or few signs of re-infection with most cells showing extensive perinuclear ER-like distribution of bacteria (Figure 6D).

**Figure 6.**
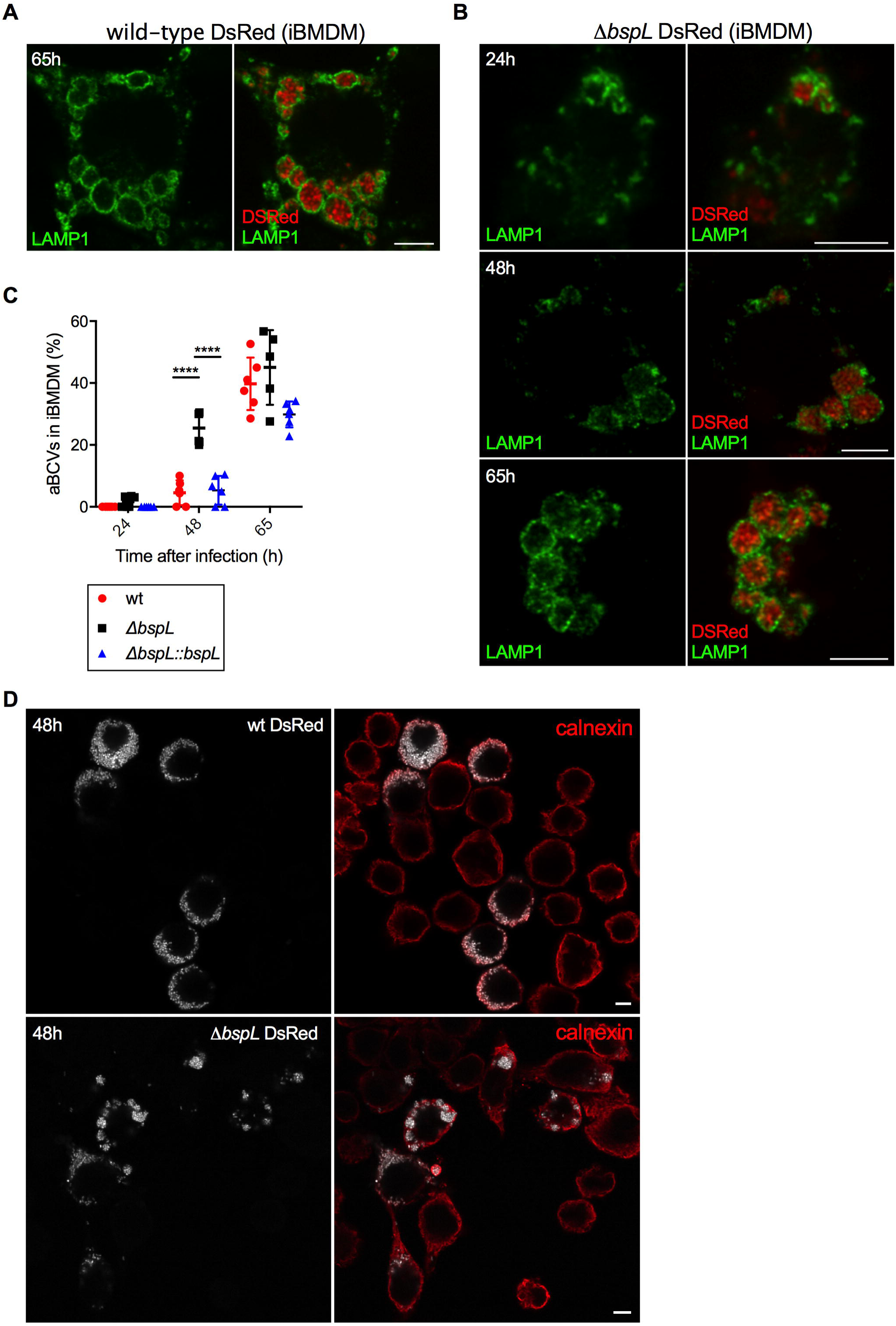
BspL is implicated in delay of aBCV formation. (A) Representative confocal images of iBMDM infected with wild-type DSred for 65h labelled for LAMP1 (green). Scale bar corresponds to 5 µm. (B) Representative confocal images of iBMDM infected with Δ*bspL* DSred for 24h (top), 48h (middle) and 65h (lower), labelled for LAMP1 (green). Scale bars correspond to 5 µm. (C) Quantification of the percentage of cells with aBCVs, in iBMDMs infected with either wild-type, Δ*bspL* or the complemented Δ*bspL::bspL* strains for 24, 48 or 65h. Data correspond to means ± standard deviations from at least 5 independent experiments. A two-way ANOVA was used yielding a P < 0.0001 (****) between wild-type and Δ*bspL* as well as Δ*bspL* and Δ*bspL::bspL* at 48h. Other comparisons are not indicated. (D) Representative confocal image of iBMDM infected with either wild-type DSRed or Δ*bspL* for 48h, labelled for calnexin (red). Bacteria shown in white. Scale bars correspond to 5 µm.

In conclusion, we propose that, secretion of BspL during *Brucella* infection induces ERAD to control aBCV formation and prevent premature bacterial egress from infected cells.

## Discussion

In this study, we characterize a previously unknown T4SS effector of *B. abortus* and its role in virulence. We found this effector hijacks the ERAD machinery to regulate the late stages of the *Brucella* intracellular cycle. Although many bacterial pathogens have been shown to control UPR, very little is known about the impact of ERAD, a downstream process following UPR, in the context of intracellular bacterial infections. To our knowledge there are only two examples. The obligatory intracellular pathogen *Orientia tsutsugamushi*, the cause of scrub thypus, is an auxotroph for histidine and aromatic amino acids and was shown to transiently induce UPR and block ERAD during the first 48h of infection (Rodino et al., 2017). This in turn enables release of amino acids in the cytosol, necessary for its growth (Rodino et al., 2017). The second example is *Legionella pneumophila*, that recruits the AAA ATPase Cdc48/p97 to its vacuole, that normally recognizes ubiquitinated substrates and can act as a chaperone in the context of ERAD to deliver misfolded proteins to the proteasome. Recruitment of Cdc48/p97 to the *Legionella* vacuole is necessary for intracellular replication and helps dislocate ubiquitinated proteins from the vacuolar membrane, including bacterial effectors (Dorer et al., 2006).

In the case of BspL we found it directly interacts with Herp, a component of ERAD which is induced upon UPR. Our data suggest that BspL enhances ERAD and this prompted us to further investigate the role of ERAD during *Brucella* infection. Interestingly, we found that inhibition of ERAD is beneficial during early stages of intracellular trafficking and enhances bacterial multiplication. It is possible that *Brucella* is transiently blocking ERAD during rBCV formation and initial replication, potentially *via* a specific set of effectors or a particular cellular signal yet to be identified. This could, as demonstrated for *Orientia*, release amino acids into the cytosol that would be critical for bacterial growth. Alternatively, or in parallel, a block of ERAD could potentially enhance autophagy to deal with the ER stress that would in turn favour rBCV formation.

As a permanent block of ERAD could become damaging to the cell under prolonged stress and, as we observed, speed up the bacterial release from infected cells potentially prematurely, *Brucella* translocation of BspL could counteract these effects by enhancing ERAD and slowing down aBCV formation. We could not directly show BspL ERAD induction is dependent on Herp as its depletion would itself block ERAD (Hori et al., 2004; Okuda-Shimizu and Hendershot, 2007). However, in the presence of BspL no glycosylated ER loaded HA-TCRα was observed indicative of enhanced processing through the ERAD pathway. Instead, only a truncated unglycosylated TCRα intermediate was detected, which disappeared in the presence of cycloheximide suggesting it is efficiently degraded. These likely correspond to a backlog of peptides awaiting proteasomal degradation, generated by an abnormal ERAD flux induced by BspL.

Further work is now required to establish the precise mechanisms that enables BspL to facilitate ERAD. It is possible that BspL interaction with Herp stabilizes it, preventing its degradation and would therefore help sustain ERAD. Indeed, ER stress significantly induces Herp levels but Herp was shown to be quickly degraded, enabling efficient modulation of ERQC (Yan et al., 2014). Alternatively, BspL may favour Herp accumulation at ERQC sites that would also enhance its ability to assist protein retrotranslocation and delivery to proteasomes. Imaging of BspL during infection will help to determine if a particular sub-ER compartment is targeted, such as ERQC-sites.

This study focuses on BspL-Herp interactions, nevertheless we cannot exclude the participation of other potential targets identified in the yeast-two hybrid screen, notably Ubiquilin 2 and Bag6. Ubiquilins function as adaptor proteins between the proteasome and ubiquination machinery and therefore participate in ERAD. Ubiquilins also interact with Herp (Kim et al., 2008) and very interestingly have been shown to play a role in control of autophagy (Şentürk et al., 2019). Our co-immunoprecipitation experiment did not reveal any binding but perhaps a weak or transient interaction is taking place not detectable with our current *in vitro* conditions. Another interesting target is Bag6, (also known as Bat3) a chaperone of the Hsp70 family that is also involved in delivery of proteins to the ER or when they are not properly folded to the proteasome. Bag6 was shown to be the target of the *Orientia* Ank4 effector that blocks ERAD (Rodino et al., 2017) and to be targeted by multiple *Legionella* effectors to control host cell ubiquitination processes (Ensminger and Isberg, 2010). Therefore, it is possible that Bag6 may contribute towards BspL control of ERAD functions during *Brucella* infection.

In addition to ERAD, we found that BspL itself was implicated in induction of UPR. However, this phenotype was independent of Herp and may be an indirect effect due to its ER accumulation or *via* another cellular target yet to be characterized. Furthermore, the increased ERAD activity upon BspL expression was not a result of increased ER stress; suggesting that BspL is independently controlling these two pathways. There is growing evidence that the induction of IRE1-dependent UPR by multiple effectors is linked to modulation of *Brucella* intracellular trafficking and intracellular multiplication (Smith et al., 2013; Taguchi et al., 2015). Our data allow us to add another piece to this complex puzzle, and place for the first time the ERAD pathway at the centre of *Brucella* regulation of its intracellular trafficking. Further work is now required to decipher all the molecular players involved.

In conclusion, our results show that ERAD modulation by BspL enables *Brucella* to temporarily delay the formation of aBCVs and avoid premature egress from infected cells, highlighting a new mechanism for fine-tuning of bacterial pathogen intracellular trafficking.

## Acknowledgements

This work was funded by the ERA-Net Pathogenomics *CELLPATH* grant (ANR 2010-PATH-006), the FINOVI foundation under a Young Researcher Starting Grant and the ANR *charm-Ed* (grant n° ANR-18-CE15-0003), both obtained by SPS. JBL was supported by a doctoral contract from the Région Rhônes-Alpes ARC1 Santé. SPS is supported by an INSERM staff scientist contract. We are very grateful to Linda Hendershot (St Judes Medical School, USA) for sending us the pcDNA-TCRα and for all the help with setting up the ERAD assay and discussion of the results. We thank Renée Tsolis (University of California at Davis, USA) and Jean Celli (Washington State University, USA) with advice for the construction of the following plasmids TEM1-VceC, HA-VceC, HA-BspB and HA-BspD, as the French Agency ANSM has prevented us from importing these vectors directly from them due to the size of the genes encoded. We also thank Thomas Henry (CIRI, Lyon, France) for the iBMDM. A final special thanks to Jean Celli (Washington State University, USA) for sending us several protocols and vectors (pSEAP and pmini-Tn7 vectors) as well as providing us constant guidance for the SEAP assay, complementation and observation of aBCVs. The two-hybrid screening was hosted by the Marseille Proteomics platform (JPB, FL) supported by Institut Paoli-Calmettes, IBISA (Infrastructures Biologie Santé et Agronomie), Aix-Marseille University, Canceropôle PACA and the Région Sud Provence-Alpes-Côte d’Azur. JPB is a scholar of Institut Universitaire de France. We thank Steve Garvis, Amandine Blanco and Arthur Louche for critical reading of the manuscript.

## Author contributions

Conceptualization: JBL, JPB, JPG and SPS. Investigation: JBL, JR, TLSL, MB, FL, KW and SPS; Writing of Original Draft: JBL and SPS; Writing, Review & Editing: all authors; Funding Acquisition: SPS.

## Supplementary Figure Legends

**Figure S1. BspL targets the ER independently of its CAAX motif without impacting ER secretion.**

(A) Schematic diagram of BspL and its domains, namely the Sec secretion signal, hydrophobic region, Prolin-rich region (PRR) and potential CAAX motif with amino acid C, T, A and N.

(B) Representative confocal images of HeLa cells expressing myc-BspL (top panel) or myc-BspLΔCAAX (bottom panel) labelled for the ER marker calnexin (red). Scale bars correspond to 5 µm.

(C) HeLa cells were co-transfected with GFP-BspL (green) and myc-BspLΔCAAX (cyan) for 24h and labelled for the ER marker calnexin (red). Scale bars correspond to 5 µm.

(D) Quantification of SEAP secretion in HEK 293T cells expressing either control empty vector (pcDNA3.1), dominant negative form of Arf1 (HA-ARF[T31N]), HA-BspL, HA, BspB or HA-BspD. Measurements were done at 24h after transfection and the secretion index corresponds to means ± standard deviations. Kruskal-Wallis with Dunn’s multiple comparisons test was used and P = 0.0164 between pcDNA control and HA-ARF[T31N] (*) and 0.0005 between pcDNA and HA-BspB (***). All other comparisons ranked non-significant.

**Figure S2. Equivalent intracellular trafficking of wild-type and *bspL* mutant strains.**

(A) Bacterial counts using colony forming units (CFU) at 2, 24 and 48h post-infection with either the wild-type (red) or Δ*bspL* strains (black) of iBMDM or (B) HeLa cells. Data correspond to means ± standard deviations from 3 independent experiments.

(C) iBMDM or (D) HeLa cells were infected with wild-type *B. abortus* DSRed (red) for 24 or 48h and labelled for the ER marker calnexin (green). Zoomed insets are indicated. Scale bars correspond to 5 µm.

(E) Quantification of the percentage of BCVs positive for calnexin or (F) LAMP1 at 24 or 48h post-infection of iBMDM with either wild-type or Δ*bspL* DSRed-expressing strains. Data are presented as means ± standard deviations from at 6 independent experiments. Kruskal-Wallis with Dunn’s multiple comparisons test was used and all comparisons between the wild-type and the mutant strain yielded P > 0.05, considered as non-significant.

**Figure S3. BspL induction of ER stress is independent of Herp.**

(A) Quantification of mRNA levels of *HERP* by quantitative RT-PCR obtained from iBMDMs infected with wild-type, Δ*bspL* or the complemented Δ*bspL::bspL* strains for 48h. Mock-infected cells were included as a negative control. Data correspond to the fold increase in relation to an internal control with non-infected cells. Data are presented as means ± standard deviations from 3 independent experiments. Kruskal-Wallis with Dunn’s multiple comparisons test was used and yielded non-significant differences.

(B) Western blot of cell lysates from HeLa cells treated with siRNA control (siCtrl) or siRNA Herp (siHerp) for 48h. A sample from non-treated cells was included as a negative control. Membrane was probed with an anti-Herp antibody followed by anti-actin for loading control.

(C) Quantification of mRNA levels of *XBP1s* or (D) *CHOP* by quantitative RT-PCR obtained from HeLa cells expressing HA-BspL or HA-VceC for 24h. Where indicated, HeLa cells were treated with siRNA control (siCtrl) or siRNA Herp (siHerp). Cells transfected with empty vector pcDNA3.1 were included as a negative control and cells treated tunicamycin at 1µg/µl for 6h as a positive control. Data correspond to the fold increase in relation to an internal control with non-transfected cells. Data are presented as means ± standard deviations from at least 3 independent experiments. Kruskal-Wallis with Dunn’s multiple comparisons test was used and yielded P=0.0184 (*) between negative siCtrl and BspL siCtrl, 0.0277 (*) between negative siHerp and BspL siHerp and 0.0485 (*) between negative siCtrl and tunicamycin siCtrl. No significant differences for observed for *CHOP*.

(E) Quantification of mRNA levels of *XBP1s* or (F) *CHOP* by quantitative RT-PCR obtained from HeLa cells expressing HA-BspL, HA-VceC or HA-BspB for 24h. Were indicated, cells were treated with 0.5 nM of TUDCA for 22h. Cells transfected with empty vector pcDNA3.1 were included as a negative control and cells treated tunicamycin at 1µg/µl for 6h as a positive control. Data correspond to the fold increase in relation to an internal control with non-transfected cells. Data are presented as means ± standard deviations from 3 independent experiments. Kruskal-Wallis with Dunn’s multiple comparisons test was used and yielded P=0.0439 (*) between BspL and BspL+TUDCA. For *CHOP*, P=0.0012 (**) between tunicamycin and tunicamycin+TUDCA, 0.0036 (**) between BspL and BspL+TUDCA and 0.0192 (*) between VceC and VceC+TUDCA. Not all comparisons are indicated.

**Figure S4. BspL induction of ERAD is ER stress-independent.**

HEK 293T cells were transfected with HA-TCRα in the absence or presence of myc-BspL for 24h. Where indicated, cells were treated with 50 µg/ml cycloheximide for the last 6h or 0.5 nM of TUDCA for 22h. The blot was probed first with an anti-TCR antibody followed by anti-actin. The same samples were loaded onto a separate (separated by dashed line) for probing with an anti-myc and anti-actin to confirm the expression of myc-BspL. Molecular weights are indicated (KDa) and relevant bands described in the text highlighted with different coloured arrows.

## Material and methods

### Cell culture

HeLa, RAW and HEK293T cells obtained from ATCC were grown in DMEM supplemented with 10% of fetal calf serum. Immortalized bone marrow-derived macrophages from C57BL/6J mice were obtained from Thomas Henry (CIRI, Lyon, France) and were maintained in DMEM supplemented with 10% FCS and 10% spent medium from L929 cells that supplies MC-CSF.

### Transfections and siRNA

All cells were transiently transfected using Torpedo® (Ibidi-Invitrogen) for 24 h, according to manufacturer’s instructions. siRNA experiments were done with Lipofectamine® RNAiMAX Reagent (Invitrogen) according the protocol of the manufacturers. Importantly, siRNA depletion of Herp was done by treatment with 3μM siRNA the day after seeding of cells and again at 24h. Depletion was achieved after 48h total. Depletion was confirmed by western blotting with an antibody against Herp. ON-TARGETplus siRNA SMARTpool (L-020918) were used for Herp and for the control ON-TARGETplus Non-targeting pool (D-001810) both from from Dharmacon. For both transfections and siRNA cells were weeded 18h before at 2×10^4^ cells/well and 1×10^5^ cells/well for 24 and 6 well plates, respectively.

### Bacterial strains and growth conditions

*Brucella abortus* 2308 was used in this study. Wild-type and derived strains were routinely cultured in liquid tryptic soy broth and agar. 50 μg/ml kanamycin was added for cultures of DSRed or complemented strains.

### Construction of BspL eukaryotic expression vectors

The BspL constructs were obtained by cloning in the gateway pDONR^TM^ (Life Technologies) and then cloned in the pENTRY Myc, HA or GFP vectors. The following primers were used 5’-GGGGACAAGTTTGTACAAAAAAGCAGGCTTCAATCGATTTTTGAAGATCACTAT-3’ and 5’-GGGGACCACTTTGTACAAGAAAGCTGGGTCCTAGTTGGCCGTGCAGAAATG-3’. For the construct without CAAX the following reverse primer was used: 5’-GGGGACCACTTTGTACAAGAAAGCTGGGTCCTAGAAATGGTCGCGACCGTCA-3’. The final constructs were verified by sequencing and expression of tagged-BspL verified by western blotting.

### Construction of *bspL* mutant and complementing strain

*B. abortus 2308* knockout mutant Δ*bspL* was generated by allelic replacement. Briefly, upstream and downstream regions of about 750 bp flanking the *bspL* gene were amplified by PCR (Q5 NEB) from *B. abortus* 2308 genomic DNA using the following primers: (i) SpeI_Upstream_Forward: actagtATGTCGAGAACTGCCTGC, (ii) BamHI_XbaI_Upstream_Reverse: CGGGATCCCGGCTC TAGAGCGCGGCTCCGATTAAAACAG, (iii) BamHI_XbaI_Downstream_Forward: CGGGATCC CGGCTCTAGAGCACCGAACCGATCAACCAG and (iv) SpeI_Downstream_Reverse: actagtCC CTATACCGAGTTGGAGC. A joining PCR was used to associate the two PCR products using the following primers pairs: (i) and (iv). Finally, the Δ*BspL* fragment was cloned in a SpeI digested suicide vector (pNPTS138). The acquisition of this vector by *B. abortus* after mating with conjugative S17 *Escherichia coli* was selected using the kanamycin resistance cassette of the pNPTS138 vector and the resistance of *B. abortus* to nalidixic acid. The loss of the plasmid concomitant with either deletion of a return to the wild type phenotype was then selected on sucrose, using the *sacB* counter selection marker also present on the vector. Deletant (Δ) strain was identified by diagnostic PCR using the following primers: Forward: CACTGGCAATGATCAGTTCC and Reverse: CTGACCATTATGTGTGAACAGG (Amplicon length: WT-2000 bp, Δ-1500 bp). The complementing strain was constructed by amplifying BspL and its promoter region (500 bp upstream) with the PrimeStar DNA polymerase (Takara) using the following primers: Fw: AAAGGATCCGACAATCAGAAGGTTTCCTATGAAACG and Rev: AAAACTAGTTCAGTTGGCCGTGCAGAAATG. Insert and pmini-Tn7 (Myeni et al., 2013) were digested with BamHI and SpeI and ligated overnight. Transformants were selected on kanamycin 50 μg/mL and verified by PCR and sequencing. To obtain the complementing strain the Δ*bspL* mutant was electroporated with pmini-Tn7-*bspL* with the helper plasmid pTNS2. Electroporants were selected on tryptic soy agar plates with kanamycin 50 μg/mL and verified by PCR.

### HA-TCR**α**

The pcDNA-TCRα was obtained from Linda Hendershort (St Judes Medical School, USA) and it corresponds to the A6-TCRα (Feige and Hendershot, 2013). The HA tag was introduced by sequence and ligation independent cloning (SLIC) method with the following primers: TCR-Fw: CGAGCTCGGATCCACTAGTCCAGTGTGGTGGAATTCTACCCATACGATGTTCCAGATTACGCTATGGGCATGATCAGCCTG and TCR-Rv:GAGCGGCCGCCACTGTGCTGGATATCTGCAGAATTCTTACTAGCTAGACCACAG. Briefly, pcDNA-TCRα was digested with EcoRI and incubated with purified PCR product amplified with the PrimeStar DNA polymerase (Takara – Ozyme) for 3 min at RT followed by 10 min on ice. The following ratio was used for the reaction: 100 ng vector + 3x PCR insert.

### Infections

Bacterial cultures were incubated for 16h from isolated colonies in TSB shaking overnight at 37 °C. Culture optical density was controlled at 600 nm. Bacterial cultures diluted to obtain the appropriate multiplicity of infection (MOI) for HeLa 1:500 and iBMDMs 1:300 in the appropriate medium. Infected cells were centrifuged at 400 x g for 10 minutes to initiate bacterial-cell contact followed by incubation for 1h at 37°C and 5% CO_2_ for HeLa cells and only 15 min for iBMDMs. After the cells were washed 3 times with DMEM and treated with gentamycin (50 μg/mL) to kill extracellular bacteria for 1h. At 2 hours pi the medium was replaced with a weaker gentamycin concentration 10 μg/mL. Cells are plated 18h before infection and seeded at 2×10^4^ cell / well and 1×10^5^cells/well for 24 and 6 well plates respectively. For qRT-PCR experiments, 10 mm cell culture plates were used at a density of 1×10^6^cell/plate. At the different time points cells were either harvested of coverslips fixed for immunostaining. In the case of bacterial cell counts, cells were lysed in 0.1% Triton for 5 min and a serial dilution plated for enumeration of bacterial colony forming units (CFU).

### Immunofluorescence microscopy

At the appropriate time point, coverslips were washed twice with PBS, fixed with AntigenFix (MicromMicrotech France) for 15 minutes and then washed again 4 times with PBS. For ER and Herp immunostaining, permeabilization was carried out with a solution of PBS containing 0.5% saponin for 30 minutes followed by blocking also for 30 minutes in a solution of PBS containing 1% bovine serum albumin (BSA), 10% horse serum, 0.5% saponin, 0.1% Tween and 0.3 M glycine. Coverslips were then incubated for 3h at room temperature or at 4 °C overnight with primary antibody diluted in the blocking solution. Subsequently, the coverslips were washed twice in PBS containing 0.05% saponin and incubated for 2h with secondary antibodies. Finally, coverslips were washed twice in PBS with 0.05 % saponin, once in PBS and once in ultrapure water. Lastly, they were mounted on a slide with ProLongGold (Life Technologies). The coverslips were visualized with a Confocal Zeiss inverted laser-scanning microscope LSM800 and analyzed using ImageJ software. For Lamp1 immunostaining no pre-permeabilization and blocking were done and coverslips were directly incubated with antibody mix diluted in PBS containing 10% horse serum and 0.5% saponin for 3h at room temperature. The remaining of the protocol was the same as described above.

### Western blotting

Cells were washed 1x with PBS and the 1x with ice-cold PBS. Cells were scrapped ince-cold PBS, centrifuged for 5 min at 4 °C at 80 g. Pellets where then ressuspended in cell lysis buffer (Chromotek) supplemented with phenylmethylsulfonyl fluoride (PMSF) and proteinase inhibitors tablet cocktail (complete Mini, Roche). Samples resolved on SDS-PAGE and transferred onto PVDF membrane Immobilon-P (Millipore) using a standard liquid transfer protocol. Membranes were blocked using PBS with 0.1% Tween 20 and 5% skim milk for 30 min and the probed using relevant primary antibodies overnight at 4 °C, washed 3 times with PBS with 0.1% Tween 20 and then incubated with HRP-conjugated secondary anti-goat, mouse or rabbit antibodies, diluted in PBS with Tween 20 0.1% and 5% skim milk for 1 h. Western blots were revealed using ECL Clarity reagent (BioRad). Signals were acquired using a Fusion Camera and assembled for presentation using Image J.

### TEM1 translocation assay

RAW cells were seeded in a 96 well plates at 1×10^4^ cells/well overnight. Cells were then infected with an MOI of 300 by centrifugation at 4 °C, 400 g for 5 min and 1 at 37 °C 5% CO_2_. Cells were washed with HBSS containing 2.5 mM probenicid. Then 6 µl of CCF2 mix (as described in the Life Technologies protocol) and 2.5 mM probenicid were added to each well, and incubated for 1.5 h at room temperature in the dark. Cells were finally washed with PBS, fixed using Antigenfix and analysed immediately by confocal microscopy (Zeiss LSM800).

### RNA isolation and real-time quantitative polymerase chain reaction (qRT-PCR)

HeLa cells were seeded in 100×100 culture dishes at 1×10^6^ cells/plate for each condition and were either transfected with HA-tagged BspL, VceC or BspB for 24h or infected with wild-type, mutant or complemented strains for 48h. Cells were then washed 1x in PBS, scrapped in buffer RLT (Qiagen) supplemented with ß-mercaptoethanol and transfered on a Qiashredder column (Qiagen). Then several wash steps were performed and total RNAs were extracted using a RNeasy Mini Kit (Qiagen). 500 ng of RNA were reverse transcribed in a final volume of 20 µl using QuantiTect Reverse Transcription Kit (Qiagen). Real-time PCR was performed using SYBR Green PowerUp (ThermoScientific) with an QuantiTect Studio 3 (ThermoScientific). Specific primers for human cells: *HERP* fw: CGTTGTTATGTACCTGCATC and *HERP* rev: TCAGGAGGAGGACCATCATTT; *XBP1s* fw: TGCTGAGTCCGCAGCAGGTG and *XBP1s* rev: GCTGGCAGGCTCTGGGGAAG; *CHOP* fw: GCACCTCCCAGAGCCCTCACTCTCC and *CHOP* rev: GTCTACTCCAAGCCTTCCCCCTGCG. The *HPRT*, and *GAPDH* expressions were used as internal controls for normalization and fold change calculated in relation to the negative control. Primers were *HPRT* fw: TATGGCGACCCGCAGCCCT and *HPRT* rev: CATCTCGAGCAAGACGTTCAG; *GAPDH* fw: GCCCTCAACGACCACTTTGT and *GAPDH* rev: TGGTGGTCCAGGGGTCTTAC.

For murine cells: *HERP* fw: CAACAGCAGCTTCCCAGAAT and *HERP* rev: CCGCAGTTG GAGTGTGAGT; *XBP1s* fw: GAGTCCGCAGCAGGTG and *XBP1s* rev: GTGTCAGAGTCCATGGGA; *CHOP* fw: CTGCCTTTCACCTTGGAGAC and *CHOP* rev: CGTTTCCTGGGGATGAGATA and for the internal controls for normalization primers were *18S* fw: GTAACCCGTTGAACCCCATT and *18S* rev: CCATCCAATCGGTAGTAGCG; *GAPDH* fw: TCACCACCATGGAGAAGGC and *GAPDH* rev: GCTAAGCAGTTGGTGGTGCA. Data were analyzed using Prism Graph Pad 6.

### ERAD evaluation

HEK293T cells seeded in 100 mm culture plates at 8×10^5^ cells/plate overnight and then co-transfected for 24h with Torpedo (Ibidi) with vectors encoding HA-TCR (5 µg) and myc-BspL (5 µg). Cycloheximide 50 µg/ml was added 6h before lysis. Where indicated, TUDCA was added 2h after transfection at 0.5 mM. Cells were harvested as described above (western blotting) and lysed in 200 µl of lysis buffer (Chromotek). EndoH (New England Biolabs) treatment was carried out following the manufacturers protocol for 1h at 37 °C. Sample buffer was then added (30 mM Tris-HCl pH 6.8, 1% SDS, 5% glycerol, 0.025% bromophenol blue and 1.25 ß-mercaptoethanol final concentration). Western blotting was done as described above using anti-TCR antibody. Actin levels were also analyzed as a loading control.

### Secretion assay

HEK293T cells were harvested and seeded in 6-well plates at 1×10^5^ cells/well and co-transfected with plasmids encoding *Brucella* secreted proteins (300 ng DNA) and the secreted embryonic alkaline phosphatase (SEAP) (300 ng DNA) provided by Jean Celli. Total amount of transfected DNA was maintained constant using an empty vector pcDNA 3.1 for the positive control. At 18 h post transfection, the transfection media was removed and then cells were still incubated at 37°C 5% CO_2_. Fourty-eight hours later, media containing culture supernatant (extracellular SEAP) was removed and collected. To obtain intracellular SEAP, each well was washed with PBS and then incubated with a solution of PBS-Triton X-100 0.5% for 10 minutes. An incubation of each fraction was performed at 65 °C following a centrifugation at maximum speed for 30 seconds. Then cells were incubated with a provided substrate 3-(4-methoxyspiro [1,2-dioxetane-3,2’(5’-chloro)-tricyclo(3.3.1.13,7) decane]-4-yl)phenyl phosphate (CSPD) by SEAP reporter gene assay, chemiluminescent kit (Roche Applied Science). Chemiluminescence values were obtained with the use of a TECAN at 492 nm. Data are presented as the SEAP secretion index, which is a ratio of extracellular SEAP activity to intracellular SEAP activity.

### Yeast two-hybrid

BspL was cloned into pDBa vector, using the Gateway technology, transformed into MaV203 and used as a bait to screen a human embryonic brain cDNA library (Invitrogen). Media, transactivation test, screening assay and gap repair test were performed as described (Orr-Weaver and Szostak, 1983; Thalappilly et al., 2008; Walhout and Vidal, 2001).

### Antibodies

For immunostaining for microscopy the following antibodies were used:

Rat anti-HA antibody clone 3F10 (Roche, #1867423) was used at a dilution 1/50 and mouse anti-HA (Covance, clone 16B12, #MMS-101R), at 1/500. Rabbit anti-calnexin (Abcam, #ab22595) was used at 1/250. Rabbit anti-Herp EPR9649 (Abcam, #ab150424) at 1/250. The mouse anti-myc antibody clone 9E10 (developed by Bishop, J.M.) was used at 1/1000. Rat anti-LAMP1 clone ID4B (developed by August, J.T.) was used 1/100 for mouse cells and mouse anti-LAMP1 clone H4A3 (developed by August, J.T. / Hildreth, J.E.K.) was used 1/100 for human cells. All LAMP1 and Myc antibodies were obtained from the Developmental Studies Hybridoma Bank, created by the NICHD of the NIH and maintained at the University of Iowa. Secondary anti-mouse, rabbit and rat antibodies were conjugated with Alexas-555, −488 or −647 fluorochromes all from Jackson Immunoresearch at a dilution 1/1000. Phallodin Atto-647 (Sigma, #65906) was used at a dilution of 1/1000. Dapi nuclear dye (Invitrogen) was used at a dilution of 1/1000.

For western blotting the following antibodies were used:

rabbit anti-FLAG (Sigma, #F7425) at 1/1000; rabbit anti-Alex3 (Sigma, # HPA000967) at 1/100; rabbit anti-Ubiquilin 2 (Abcam, #ab217056) at 1/1000; rabbit anti-Herp EPR9649 (Abcam, # ab150424) at 1/1000; mouse anti-HA (Covance, clone 16B12, ref. MMS-101R) at 1/1000; rabbit anti-TCR clone 3A8 (Invitrogen, #TCR1145) at 1/1000; mouse anti-myc antibody clone 9E10 at 1/1000; mouse anti-actin AC-40 (Sigma, #A4700) at 1/1000. Anti-mouse (GE Healthcare) or rabbit-HRP (Sigma) antibodies were used at 1/5000.

### Drug treatments

All drug treatments are indicated in the specific protocols. To summarize the concentrations used were: TUDCA (Focus Biomolecules) at 0.5 nM; Cycloheximide (Sigma) at 50 µg/ml; Eeyarstatin (Sigma) at 8 µM; Tunicamycin (Sigma) at 1 µg/µl; Probenicid (Sigma) at 2.5 mM.

### Co-immunoprecipitation

HeLa cells were cultured in 100 mm x 20 mm cell culture dishes at 1×10^6^ cells/dish overnight. Cells were transiently transfected with 30 uL of Torpedo ^DNA^ (Ibidi) for 24h for a total of 10 µg of DNA/plate. On ice, after 2 washes with cold PBS cells were collected with a cell scraper and centrifuged at 80g at 4 °C during 10 min. Cell lysis and processing for co-immunoprecipitation were done as described with the PierceTM HA Epitope Antibody Agarose conjugate (Thermo scientific).

### Statistical analysis

All data sets were tested for normality using Shapiro-Wilkinson test. When a normal distribution was confirmed a One-Way ANOVA test with a Tukey correction was used for statistical comparison of multiple data sets with one independent variable and a Two-Way ANOVA test for two independent variables. For data sets that did not show normality, a Kruskall-Wallis test was applied, with Dunn’s correction, or Mann-Whitney U-test for two sample comparison. All analyses were done using Prism Graph Pad 6.

