## Supplementary figures and images for "*Brucella* effector hijacks endoplasmic reticulum quality control machinery to prevent premature egress"

### Supplemental Figure 1

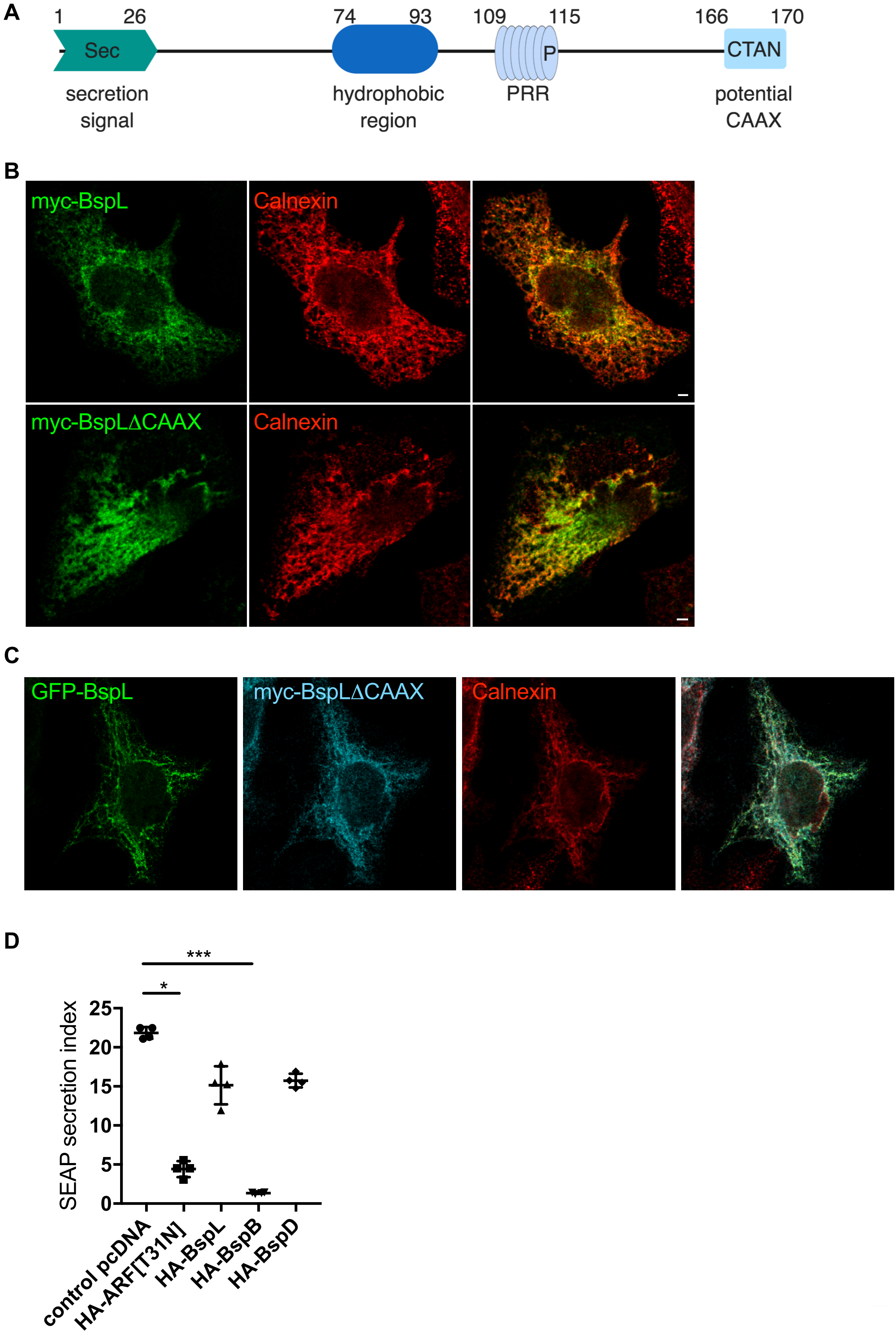

### Supplemental Figure 2

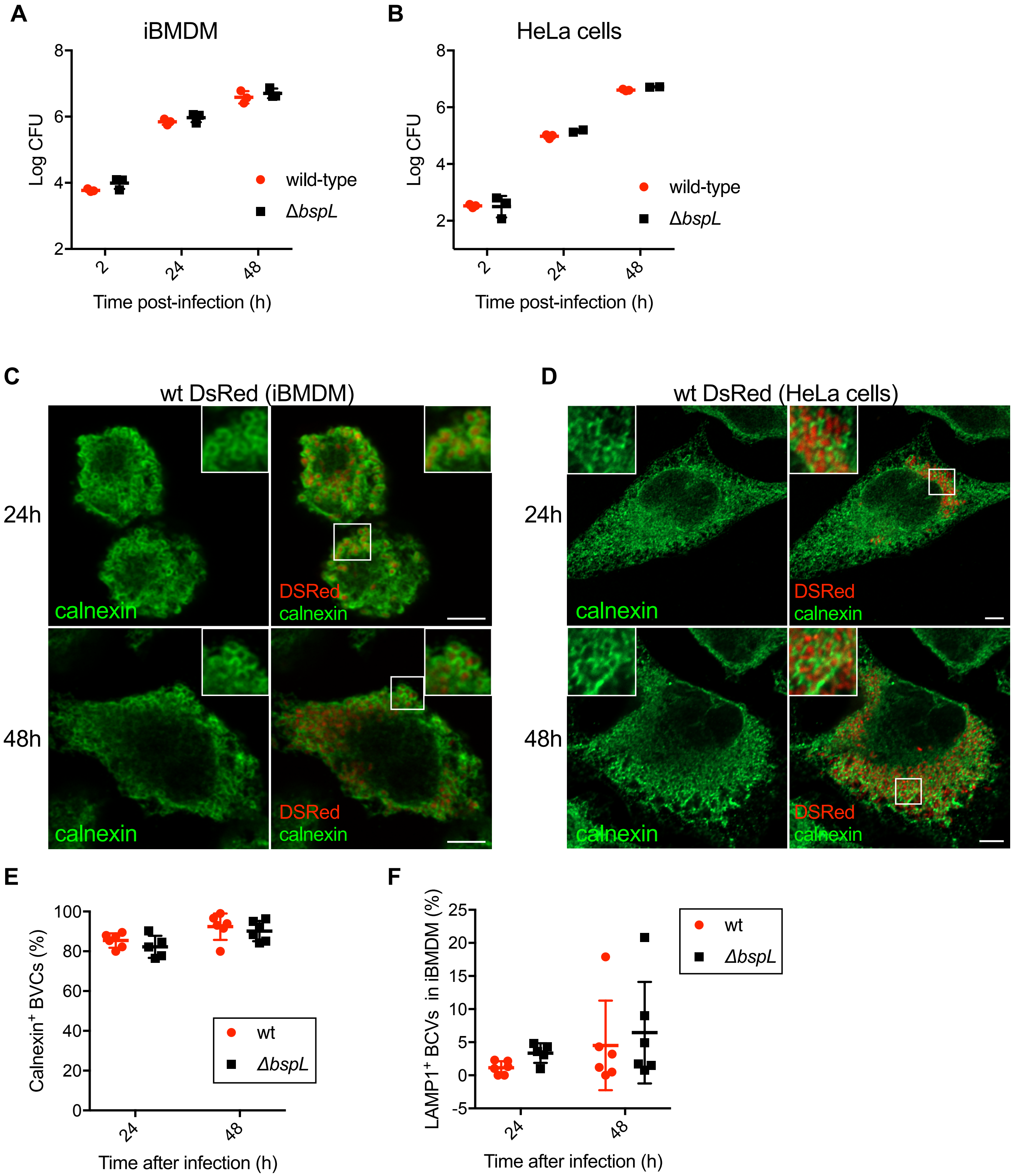

### Supplemental Figure 3

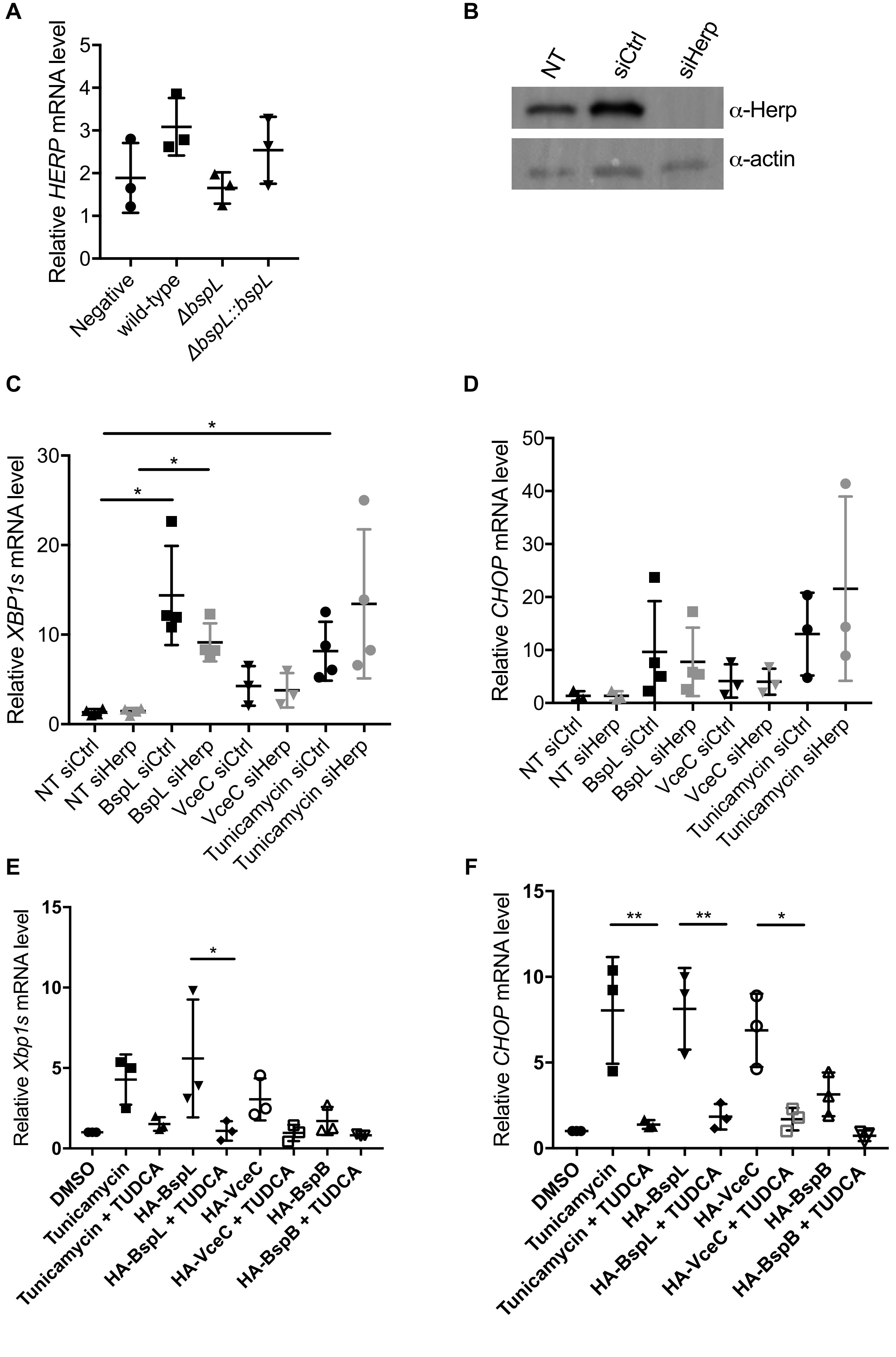

### Supplemental Figure 4

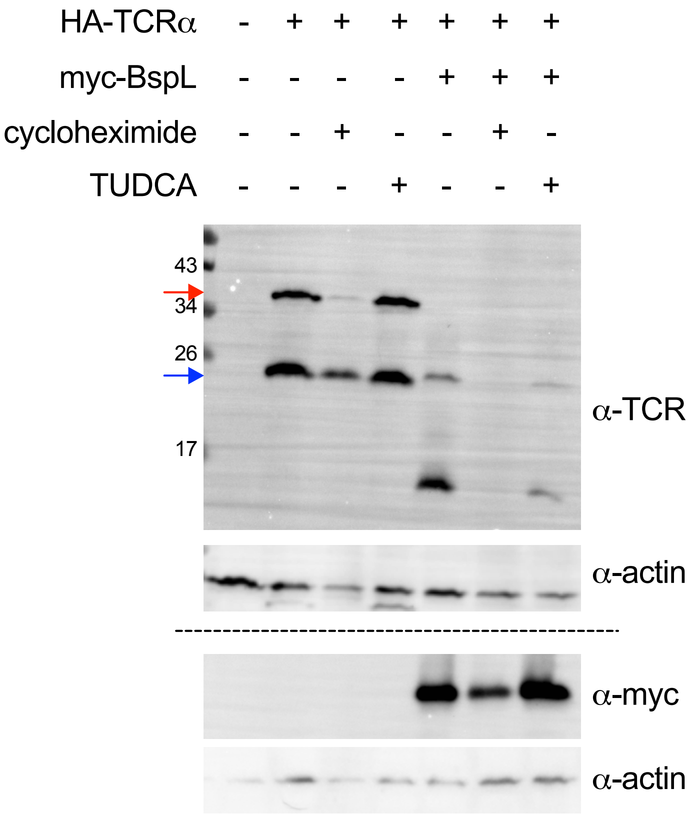
